# Base-resolution DNA methylome of human MDS hematopoietic stem cell reveals TET2-GFI1 epigenetic axis repressing MDS

**DOI:** 10.64898/2026.02.10.704995

**Authors:** Liangding Hu, Qicong Shen, Yan Gu, Jiahui Lu, Yuhang Li, Na Liu, Bin Zhang, Yanmei Han, Qian Zhang, Xuetao Cao

## Abstract

Dysfunctional hematopoietic stem cells (HSCs) drive the initiation of myelodysplastic syndromes (MDS), yet the genome-wide DNA methylation landscape in MDS primitive HSCs and its mechanistic contribution to disease pathogenesis remain poorly defined. Here, we establish single-base resolution DNA methylomes of bone marrow HSCs from MDS patients and healthy donors. We uncover widespread hypermethylation within CpG islands, alongside hypomethylation in repetitive elements such as Alu. Differentially methylated regions (DMRs) are enriched for genes involved in cancer-related pathways, as well as extrinsic signaling pathways and intrinsic transcriptional networks essential for HSC function. Among these, we identify GFI1 and BMI1 as key targets of DNA methylation dysregulation in MDS. Notably, using either MDS or Tet2-deficient mouse model, we demonstrate that loss of Tet2, a frequently mutated epigenetic regulator in MDS, induces promoter hypermethylation and transcriptional repression of Gfi1, contributing to expansion of the MDS or aged hematopoietic stem and progenitor cell pool. Our study not only charts the base-resolution DNA methylome of human MDS HSCs but also reveals a TET2-GFI1 axis that safeguards HSC homeostasis. These findings provide mechanistic insight into how aberrant DNA methylation drives HSC dysfunction in MDS and offer an epigenomic resource for discovering novel regulators and therapeutic targets at the stem cell level.

## Introduction

Myelodysplastic syndromes (MDS) represent a heterogeneous group of clonal hematopoietic disorders characterized by ineffective hematopoiesis, peripheral cytopenias and a heightened risk of progression to acute myeloid leukemia (AML)^1^. Accumulating evidence positions hematopoietic stem cells (HSCs) as the cellular origin of MDS, where both intrinsic transcriptional dysregulation and extrinsic microenvironmental signals converge to disrupt normal self-renewal and differentiation^2-4^. Among the epigenetic alterations implicated in MDS pathogenesis, DNA methylation abnormalities have emerged as a critical driver, often associated with disease progression and poor prognosis^5^. Previous studies using relative low-resolution or candidate-region approaches have reported widespread methylation changes in MDS, including hypermethylation at tumor suppressor gene promoters^6-8^. However, a whole-genome base-resolution map of DNA methylomes in human primitive MDS HSCs revealing the systemic epigenetic dysregulation at stem-cell level is still lacking.

TET2, a member of the ten-eleven translocation family of DNA dioxygenases, catalyzes the oxidation of 5-methylcytosine and plays a key role in active DNA demethylation^9^. Tet2 plays essential roles in establishing cell-specific function of myeloid cells and their progenitors^10-12^. The ten-eleven-translocation (TET1-3) proteins are α-ketoglutarate- and Fe2^+^-dependent dioxygenases that catalyze the iterative oxidation of both DNA and RNA 5-methylcytosine (5-mC) to 5-hydroxymethylcytosine (5-hmC), 5-formylcytosine (5-fC), and 5-carboxylcytosine (5-caC). These oxidized mCs are key intermediates mediating DNA demethylation through replication-dependent dilution or base excision repair (BER)^13^. Recurrent loss-of-function mutations in TET2 are frequently observed in MDS and are associated with increased self-renewal of hematopoietic stem and progenitor cells (HSPCs)^14-16^. Nevertheless, whether and how TET2 maintains the expression of key regulators, especially in the context of human MDS HSCs via DNA methylation-centered mechanisms still needs further investigation.

In this study, we performed whole-genome bisulfite sequencing (WGBS) at single-base resolution to systematically compare the DNA methylome of bone marrow primitive HSCs isolated from MDS patients with that from healthy donors. We aimed to investigate the mechanisms of MDS pathogenesis and identify the potential targets for MDS treatment by focusing on charting the patterns of aberrant DNA methylation and the dysregulated genes in MDS HSCs, and revealing how epigenetic regulator, frequently mutated or dysregulated in MDS, regulate malignant transformation at HSC level. Our work not only provides a high-resolution epigenetic resource for MDS research but also identifies GFI1 as a critical target through which TET2 maintains HSC homeostasis and suppresses MDS development. These findings advance our mechanistic understanding of epigenetic dysregulation in MDS and highlight potential avenues for stem-cell-directed therapeutic intervention.

## Results

### Establishing base-resolution DNA methylome of primitive HSC from healthy donors and MDS patients

The advanced MDS patients, i.e. high-risk refractory anemia with excess blasts (RAEB), have hypermethylation variations in several tumor-suppressor gene loci^5,17^. Hypomethylating agents have become foundation therapies in higher-risk myelodysplastic syndrome (MDS)^18,19^. We mainly focused on MDS patients of REAB-1/2 in this study. To elucidate the specific DNA methylation profiling and discover the potential pathogenic role of DNA methylation in MDS HSCs, we performed genome-wide DNA methylome at single-base resolution by using whole genome bisulfite sequencing (WGBS)^20^. In addition to CD34, we used CD133, which is a positive marker of primitive HSC^21^, to differentiated HSC from myeloid progenitors and blast. The published scRNA-seq data of bone marrow CD34^+^ cells from MDS patients also confirmed our HSC isolation strategy: CD34^+^CD133^+^ cells are negative for CD38, CD117 and CD123 which are markers of myeloid progenitors and blast^22^ **(Fig. S1**). DNA samples were extracted from lin^−^CD34^+^CD133^+^ primitive HSCs that were freshly isolated by sorting from bone marrow of healthy donors (N) and MDS patients (M) (**Table S1**). According to chromosome distribution analysis, we also discovered +8 chromosome variations in two MDS samples (**Fig. S2a**). Among the total methylcytosines (mCs) of human HSCs detected in both normal and MDS samples, about 97% mCs are in the CG context, not like human embryonic stem cell (H1 type) genome containing a higher fraction of non-CG mCs (**Fig. S2b**). Moreover, compared with that in H1 embryonic stem cells, the total level of DNA methylation in normal HSCs is lower, regardless of their sequence context (**Fig. S1c**). To reveal the genomic patterns of DNA methylation of normal HSCs, we performed analysis of DNA methylation in CG context and CG densities in both whole genome and different genomic elements. We found that, although different functional genomic regions had their own characteristic DNA methylation patterns, DNA methylation levels decreased when CG densities increased (**Fig. S3**).

### MDS HSCs possess specific DNA methylation variations in gene-associated regulatory regions

We then went further to investigate the DNA methylation difference between normal and MDS HSCs. In the genomic view, the significant difference of both proportions and methylation levels of mCs were not found between normal and MDS samples (**Fig. 1a, b and Fig. S4a**). We then investigated whether DNA methylation variation in CG context could differentiate MDS HSCs from normal ones in specific genomic elements. By clustering the mean DNA methylation level of each of the genomic elements, we segregated these MDS samples from the normal ones. In the heatmap, we could directly visualize that increased DNA methylation levels in the elements of CpG islands (CGIs), CGI shores, tRNA and 5’-UTR in at least two MDS samples, compared with those in the normal samples (**Fig. 1c**). PCA analysis of DNA methylation levels in DNA elements of 5’-UTR, CGI and tRNA, but not proximal promoters could differentiate MDS HSCs from normal HSCs, especially in 5’-UTR (**Fig. 1d and Fig. S4b**). Moreover, by analyzing DNA methylation atlas in gene-associated regions, we also observed higher mean DNA methylation levels in CG context in 5’-UTR in MDS HSCs than those in normal HSCs (**Fig. 1e**). While in non-CG context, MDS HSCs had higher DNA methylation levels across the entire gene loci than normal ones, although they both showed low DNA methylation status (**Fig. S4c**). Notably, such trends were more obvious in samples of M1 and M2, implying heterogeneous DNA methylation variations among different MDS samples, probably due to different clinical features. Through establishing base-resolution DNA methylome of human MDS HSCs, we found increased DNA methylation level in several genomic elements associated with gene expression regulation.

**Fig. 1.**
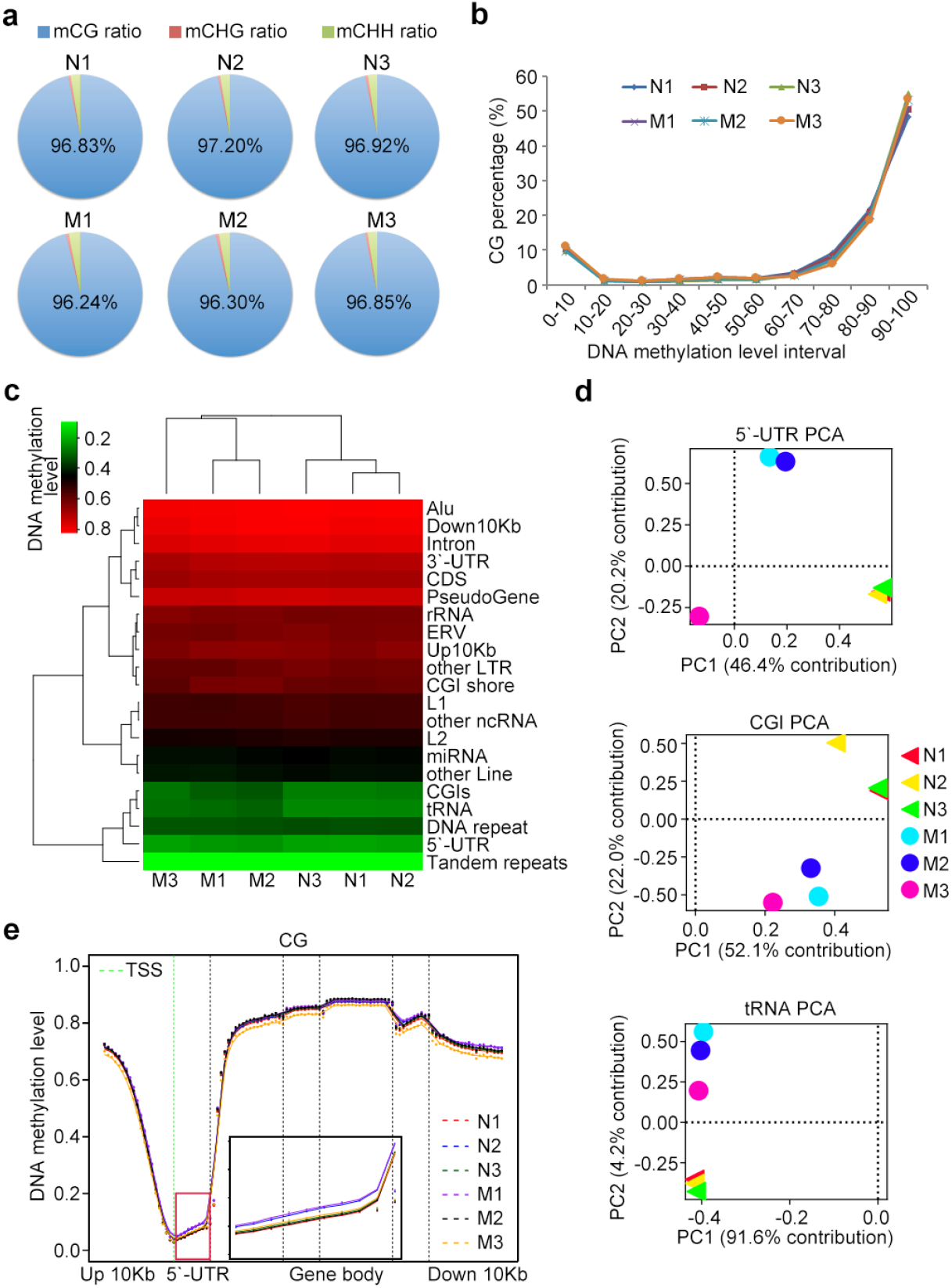
Genome-wide trends of DNA methylation of HSCs in healthy donors (N) and MDS patients (M) **a**. The proportion of methylcytosines (mCs) in CG, CHG, and CHH sequence context. H represents any nucleotide except G. **b**. Distribution of DNA methylation levels of cytosines (Cs) in CG context. **c**. Heatmap analysis of two-dimensional hierarchical clustering of mean DNA methylation levels in CG context of different genomic features. **d**. PCA analysis of DNA methylation levels in CG context of indicated genomic elements. **e**. Mean DNA methylation levels for cytosines (Cs) in CG context across all gene loci. Equal-sized bins are used for calculating mean DNA methylation level. The part for DNA methylation level of 5’-UTR was enlarged. up: upstream, down: downstream.

### CGI hypermethylation and repetitive DNA hypomethylation characterize MDS HSCs

Differentially methylated regions (DMRs) play crucial roles in regulating transcription of key genes in malignant occurrence and development^23^. Thus, we searched specific DMRs across the whole genome between the two groups (**Table S2**). DMRs in CGI, which were discovered to be the most DNA methylation aberrant regions in cancers^24^, are preferentially hypermethylated ones, while DMRs in repetitive elements were preferentially hypomethylated ones, in MDS HSCs, compared with normal HSCs (**Fig. 2a**). Notably, DMRs in 5’-UTR outside CGIs analyzed here, were not preferentially hypermethylated ones in MDS HSCs. Together with the data in **Figure 1**, these results indicate increased DNA methylation levels in 5’-UTR-located CGIs in MDS HSCs. We then separately analyzed the two kinds of DMRs and validated increased mean DNA methylation levels across CGI-located and hypermethylated DMRs in MDS samples especially M1 and M2 in profiling analysis (**Fig. 2b**). Moreover, total DMRs or CGI-located DMRs with increased DNA methylation levels could differentiate MDS HSCs from normal ones, although heterogenous DNA methylation variations were also observed among MDS samples (**Fig. 2c**). Increased DNA methylation levels in hypermethylated DMRs could also be directly visualized in DMR-centered clustering assay in MDS samples, which also showed less increased DNA methylation levels in M3 compared with the other two (**Fig. 2d**).

**Fig. 2.**
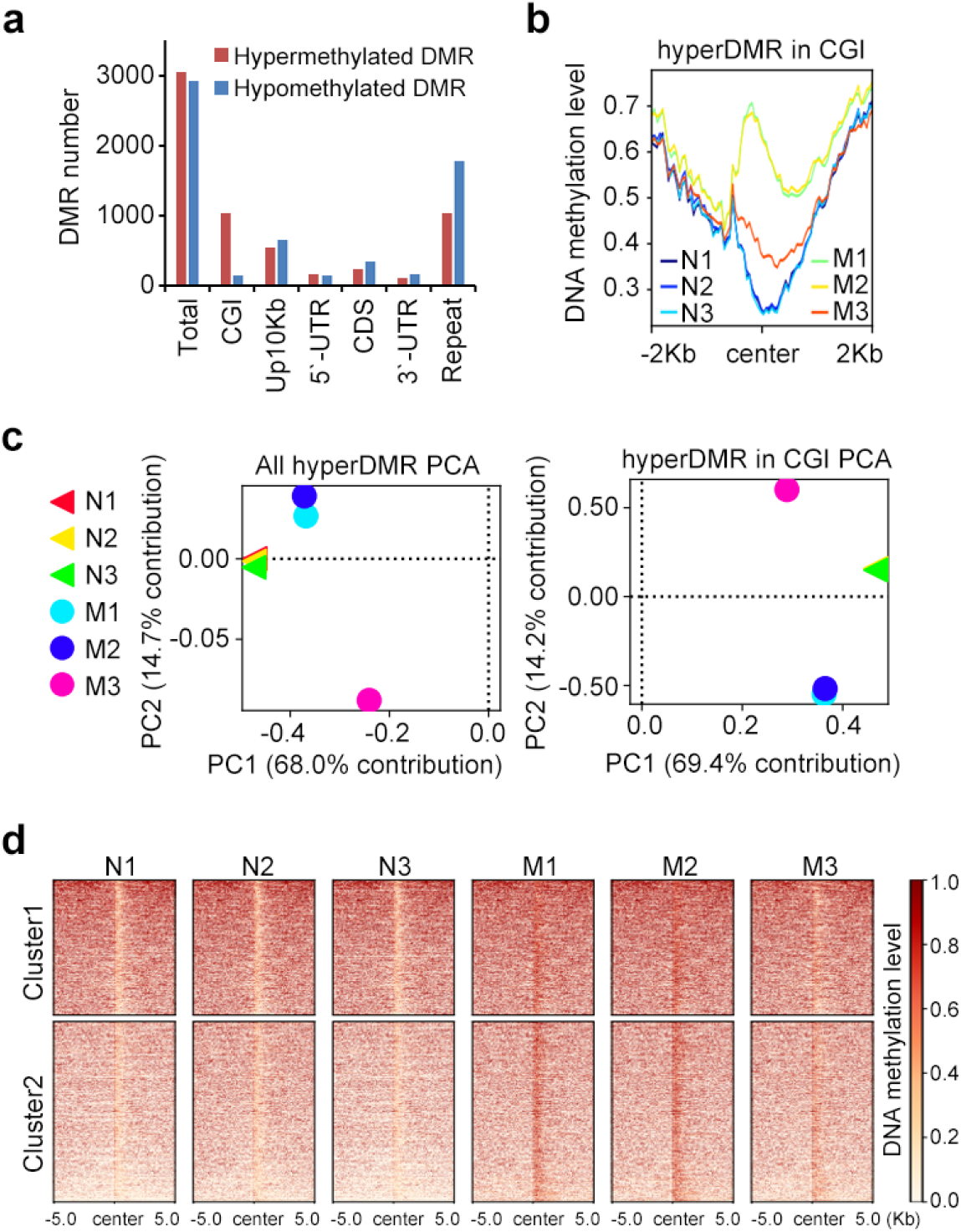
Genomic patterns with increased DNA methylation levels in MDS HSCs. **a**. Distributions of DMRs across different genomic elements. Intragenic regions analyzed here were outside CGIs. **b**. Profiling analysis of mean DNA methylation levels across CGI-located hypermethylated DMRs (hyperDMR) in MDS HSCs compared with normal ones. **c**. PCA analysis of DNA methylation levels in total or CGI-located hypermethylated DMRs. **d**. Clustering analysis of DNA methylation levels across hypermethylated DMRs.

For DMRs with downregulated DNA methylation levels, we found they were enriched in several kinds of repetitive DNA elements especially in Alu (**Fig. 3a**). We validated decreased mean DNA methylation levels across Alu-located and hypomethylated DMRs in MDS samples especially M1 and M3 in profiling analysis (**Fig. 3b**). Moreover, total DMRs or Alu-located DMRs with decreased DNA methylation levels could differentiate MDS HSCs from normal ones, although heterogenous DNA methylation variations were also observed among MDS samples (**Fig. 3c**). Decreased DNA methylation variations in hypomethylated DMRs could also be directly visualized in DMR-centered clustering assay in MDS samples, which also showed less decreased DNA methylation levels in M2 compared with the other two (**Fig. 3d**). Collectively, these results imply the roles of de novo DNA methylation-associated gene silencing and DNA demethylation-associated genomic instability in dysregulation of MDS HSCs.

**Fig. 3.**
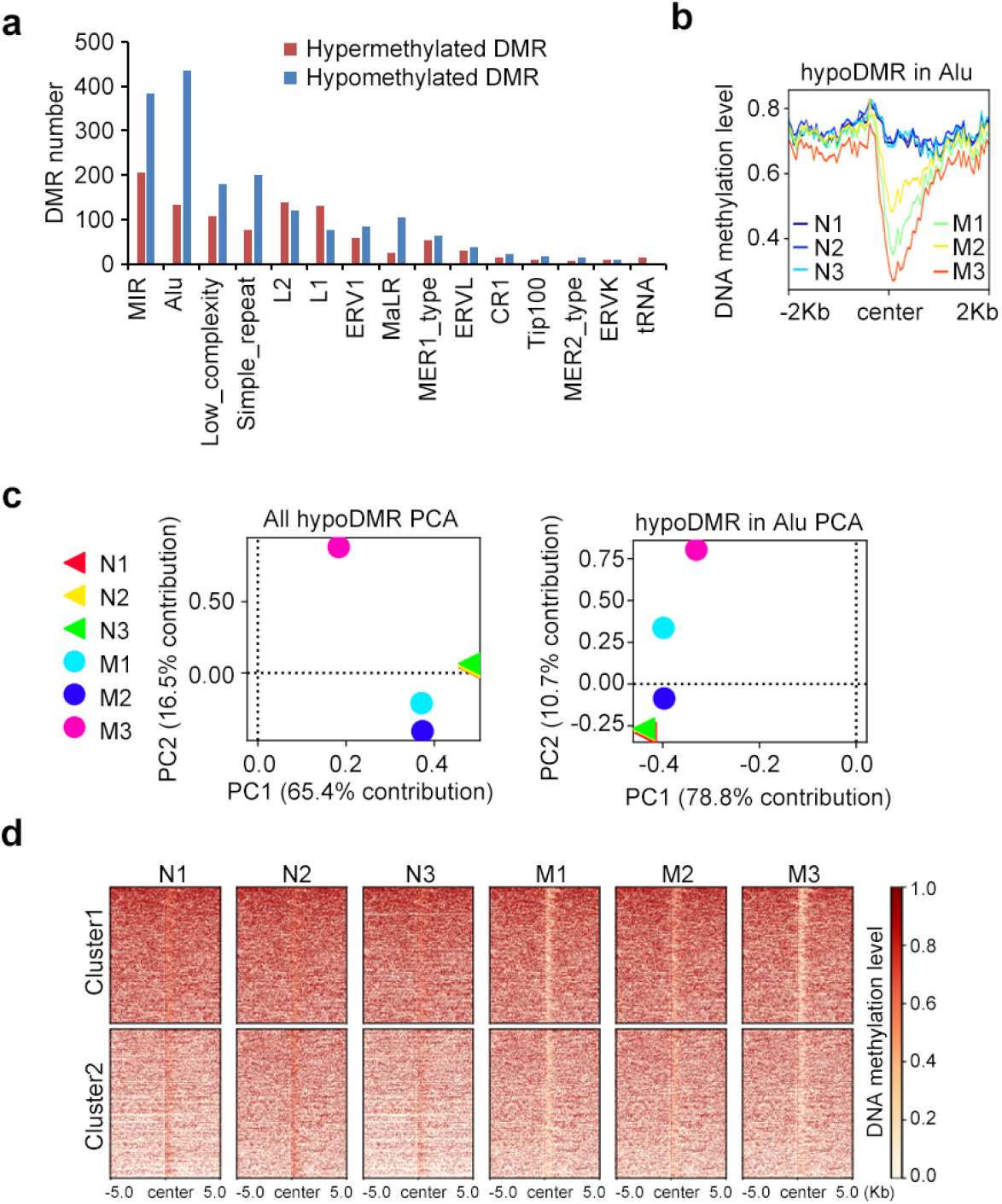
Genomic patterns with decreased DNA methylation levels in MDS HSCs. **a**. Distributions of DMRs across repetitive elements which were outside CGIs. **b**. Profiling analysis of DNA methylation levels in Alu-located hypomethylated DMRs (hypoDMR) in MDS HSCs compared with normal ones. **c**. PCA analysis of DNA methylation levels in total or Alu-located hypomethylated DMRs. **d**. Clustering analysis of DNA methylation levels across hypomethylated DMRs.

### DNA methylation variations in intrinsic and extrinsic regulators in MDS HSCs

To search for the HSC-related key genes dysregulated by aberrant DNA methylation in MDS patients, we performed pathway enrichment analysis for the DMR-associated genes (**Fig. 4a and Table S3**). Besides the cancer-related pathways that were most enriched with hypermethylated DMR-associated genes, we found that some key pathways regulating HSCs maintenance were also significantly enriched, such as Wnt and MAPK signaling pathways. For hypomethylated DMR-associated genes, pathways related to cytokines and chemokines were mostly enriched. Among these pathways, some gene loci such as IL-6, IFN-γ receptors, GM-CSF, G-CSF, IL-2 receptors exhibited hypomethylation abnormalities in MDS HSCs. These genes play important pathological functions in cancer-related inflammation, HSCs proliferation, homing and differentiation^25^. We charted these DMR-associated genes together with their DNA methylation variations (**Fig. 4b**).

**Fig. 4.**
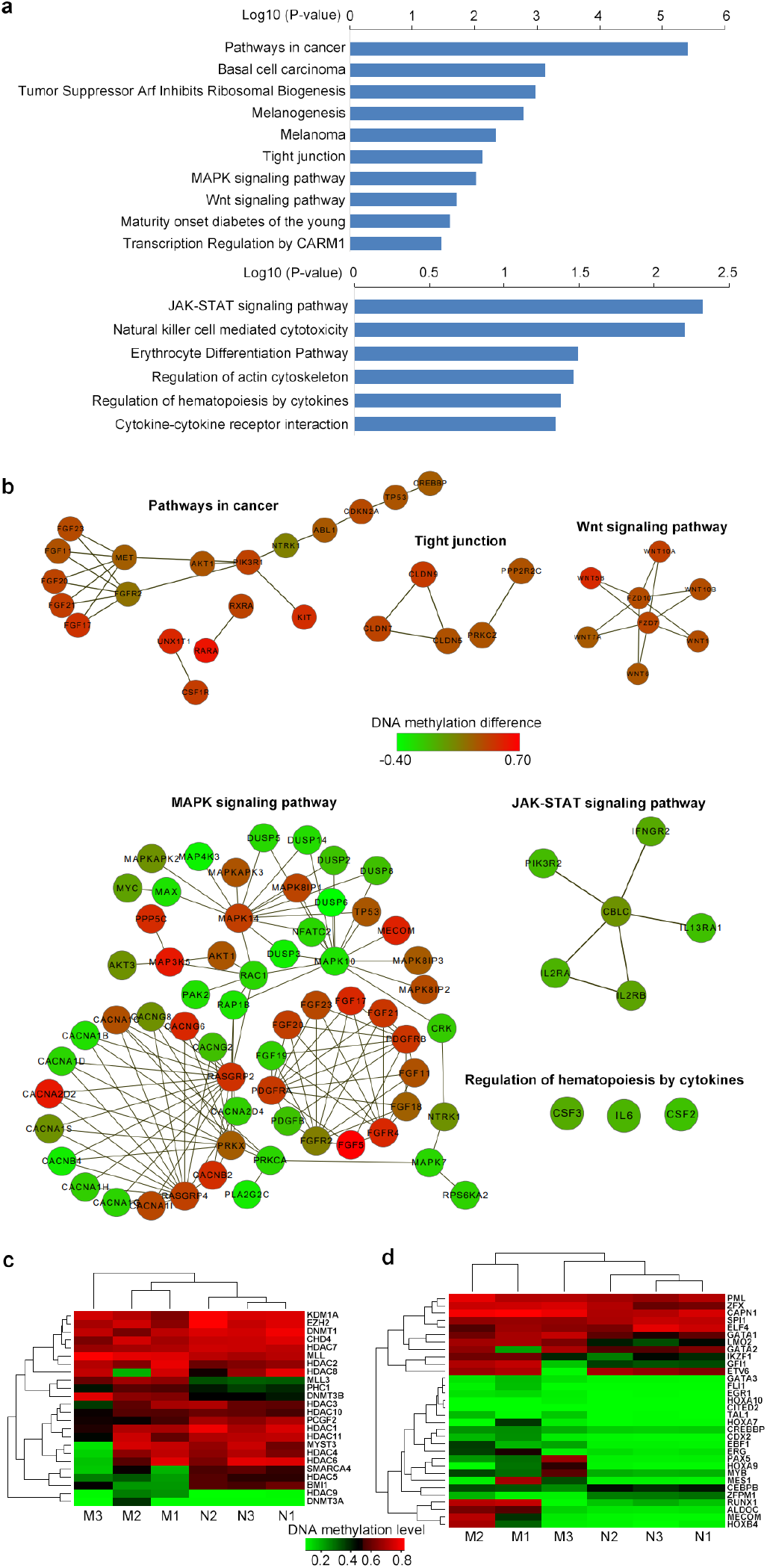
DNA methylation variations of extrinsic and intrinsic regulators in MDS HSCs. **a**. Pathways significantly enriched with DMR-associated genes (*p* < 0.05). Top panel, hypermethylated DMR; Bottom panel, hypomethylated DMR. **b**. Genes in representative pathways. The color of each circle represented the DNA methylation variation levels between normal and MDS samples in DMRs in each gene (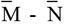, M: MDS samples included in DMR, N: Normal samples). The black line represents the interaction between two genes. **c, d**. Hierarchical clustering of DNA methylation levels of epigenetic regulators and transcription factors. The value was DNA methylation level of each gene which was determined as follows: We used 500bp window sliding from upstream 10Kb to 3’-UTR of each gene locus to find the mostly differentially methylated window between normal and MDS samples. DNA methylation level of the window for each sample was used here.

As the key intrinsic regulators of HSCs, transcription factors and epigenetic regulators are crucial in regulating HSCs self-renewal and differentiation^3^, and are also largely associated with DMR-associated genes (**Table S4**). To reveal the correlations of intrinsic regulators and aberrant DNA methylation during MDS development, we analyzed DMR-associated genes in light of previously reported key intrinsic regulators and clustered their DNA methylation variations^5,24,25^, identifying the distinct DNA methylation differences for these regulators between the two groups. We found that transcription factors including GFI1, RUNX1, CEBPB and ERG, and epigenetic regulators including BMI1, HDAC5, SMARCA4, DNMT3B, MLL3 and PHC1, are associated with DNA methylation variations at least in two MDS samples (**Fig. 4c, d**). These data imply that DNA methylation could dysregulate key regulators in gene transcription, which may lead to extensive epigenetic or transcription variations in MDS HSCs.

### Tet2 promotes Gfi1 expression via promoter demethylation to inhibit expansion of MDS HSCs

According to the data, we noticed that GFI1 and BMI1 were respectively associated with hypermethylated or hypomethylated DMRs in two MDS samples (M1 and M2). The hypermethylated DMR was across the 5’-UTR in *GFI1* gene locus, and notably, there were also DNA methylation variations among MDS samples at *GFI1* promoter (**Fig. 5a**). The hypomethylated DMR was in upstream of *BMI1* gene locus (**Fig. S5a**). To verify the universality and clinical correlation of GFI1 and BMI1 with aberrant DNA methylation, we tested another 12 HSCs samples of MDS patients and found the consistence of DNA methylation abnormalities in the other 8 samples of RAEB cases, while those of the two tAML cases and the single RA case were similar with M3 sample, further implicating the heterogeneity among different MDS subtypes. The mRNA levels of *GFI1* and *BMI1* were also dysregulated accompanied with the aberrant DNA methylation (**Fig. 5b, c and S5b, c**). Due to the small cohort size of MDS subtypes in this study and the high heterogeneity of MDS, we cannot conclude that there is no DNA methylation variation in GFI1- and BMI1-associated DMR in other MDS subtypes. However, according to our data, the two DNA methylation variations were common among patients of RAEB subtype.

**Fig. 5.**
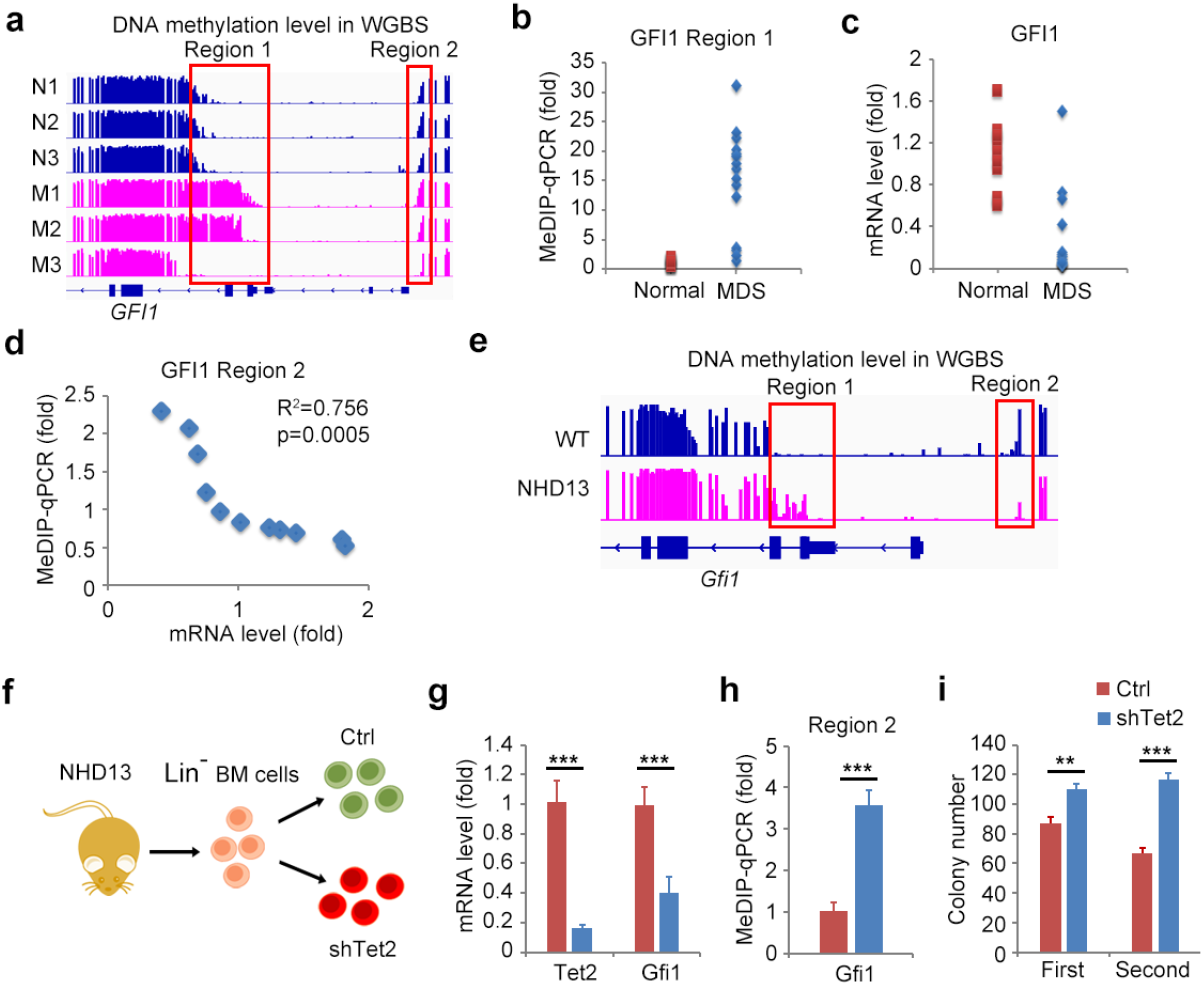
TET2 represses DNA methylation at *GFI1* promoter to restrict stem pool of MDS. **a**. DNA methylation levels in CG context across *GFI1* gene in genome browser view. Red boxes show regions with DNA methylation variations in normal (N) and MDS (M) HSCs. **b, c**. MeDIP-qPCR analysis of DNA methylation levels of region 1 in *GFI1* locus and RT-qPCR analysis of mRNA levels of GFI1 in normal (n = 10) and MDS (n = 15) HSCs. **d**. MeDIP-qPCR analysis of DNA methylation levels of region 2 in *GFI1* locus and RT-qPCR analysis of mRNA levels of GFI1 in MDS HSCs with increased DNA methylation levels in region 1 compared with normal HSCs (n = 11). **e**. DNA methylation levels across *Gfi1* gene in genome browser view (GSE129691). Red boxes show regions with DNA methylation variations in control (Ctrl) and MDS (*NHD13*) murine HSCs. **f-i**. RT-qPCR analysis of mRNA levels of indicated genes, MeDIP-qPCR analysis of DNA methylation levels in region 2, and colony formation analysis of the serial replating assay of the control and lentivirus-mediated Tet2-knockdown (shTet2) Lin^-^ HSPCs from NHD13 mice. Data were normalized by input DNA (**b, d, h**) or β-actin (**c, d, g**), and compared with normal samples or control groups. Data are the mean ± s.d. (**g**-**i**), two-tailed unpaired Student’s *t*-test. *, *P* < 0.05; **, *P* < 0.01; ***, *P* < 0.001

Interestingly, we noticed that although similar DNA methylation levels in region 1 of M1 and M2 MDS samples, the mRNA level of GFI1 in M2 sample was higher than that in M1 sample (**Fig. S5d**), implying other regulatory mechanism between M1 and M2 samples in *GFI1* gene loci. We noticed that DNA methylation level of region 2 in M2 sample was lower than that in M1 sample (**Fig. 5a**), implying that DNA methylation level of region 2 might regulate GFI1 expression in MDS HSCs. We then investigated the correlation between DNA methylation levels of region 2 and GFI1 mRNA levels in MDS samples which had higher DNA methylation levels in region 1 than those in normal samples, and found that they were negatively correlated (**Fig. 5d**), implying that DNA methylation in region 2 which locates at *GFI1* promoter could repress GFI1 expression.

Tet2 acts as a tumor repressor in myeloid cancer including MDS. In a Nup98-HoxD13 (NHD13) transgenic mouse model of MDS, Tet2 loss further expands the hematopoietic stem/progenitor pool and accelerates leukemia transformation^26^. We went further to investigate whether Tet2 could promote Gfi1 expression through mediating DNA demethylation at *Gfi1* promoter. Using public data of DNA methylome of HSPCs from NHD13 mouse^27^, we firstly investigated that, although there was increased DNA methylation level cross 5’-UTR of Gfi1 in HSCs from NHD13 mice, compared to wildtype control, decreased DNA methylation level at *Gfi1* promoter was observed (**Fig. 5e**). We then silenced Tet2 in HSPCs from NHD13 mice, and found that Tet2-knockdown (Tet2-KD) increased DNA methylation level at *Gfi1* promoter, and decreased mRNA level of Gfi1 (**Fig. 5f-h**). Colony forming cell (CFC) assays revealed that Tet2-KD in HSCs from NHD13 mice could exhibit not only a higher CFC number but also higher replating capacity, compared with control HSCs from NHD13 mice (**Fig. 5i**). These results imply that Tet2 promotes promoter demethylation to maintain high expression of GFI1 in HSCs to repress MDS development.

### Tet2 loss decreases Gfi1 expression via promoter hypermethylation for expansion of aged HSCs

As loss of function mutations of Tet2 contribute to MDS development, and aged but not young Tet2 knockout mice showed enlargement of the hematopoietic stem cell compartment and myeloproliferation^14^. We went further to validate that promoting Gfi1 expression through promoter demethylation in HSCs was a key mechanism for Tet2-mediated inhibition of MDS development. Using a published dataset of RNA-seq and WGBS analysis of young or aged bone marrow Lin^-^Sca-1^+^c-Kit^+^ stem cells (LSK) from Tet2 catalytic mutant (Mut) and knockout (KO) mice^28^, we analyzed genes with decreased mRNA levels due to Tet2 loss in aged but not young LSK. Interestingly, we found that a large fraction of these genes showed increased mRNA levels in aged wild-type control LSK (**Fig. 6a**), and these genes were enriched in cancer-associated and neural signaling pathways (**Fig. 6b and S6**), implying that impaired transcription increase of these genes in aged HSCs due to Tet2 loss may contribute to HSC transformation, which was investigated in aged Tet2-KO mice. Notably, Gfi1 was in the list (**Fig. 6c**). We further investigated the role of Tet2-repressed DNA methylation in regulating expression of Gfi1. Interestingly, both bone marrow LSK and Lin^-^ progenitor cells from both Tet2-Mut and Tet2-KO young mice showed slightly increased DNA methylation levels at *Gfi1* promoter (**Fig. 6d**), further implying Tet2-mediated DNA demethylation in promoting Gfi1 expression against HSC Aging.

**Fig. 6.**
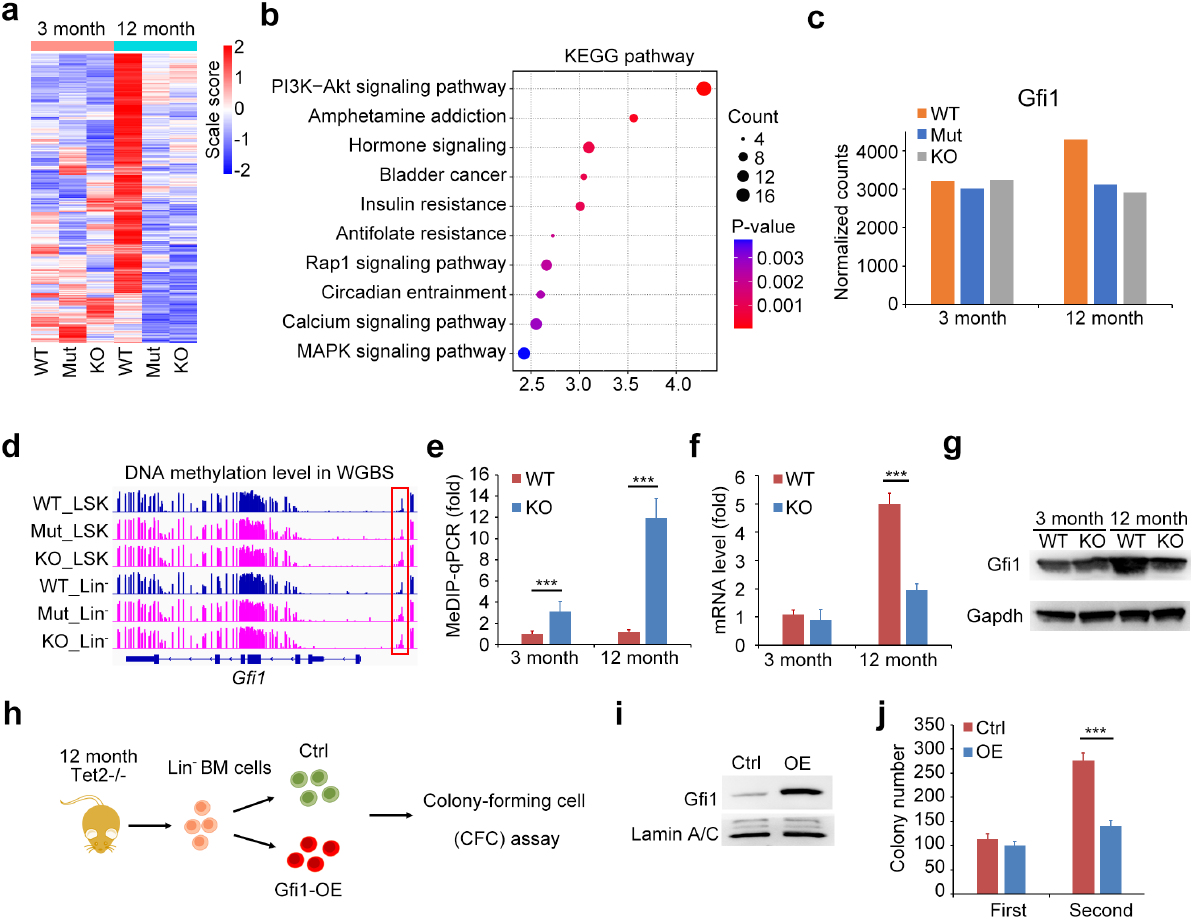
Loss of Tet2 leads to DNA hypermethylation-mediated repression of Gfi1 expression in aged HSCs. **a-c**. Heatmap presentation and KEGG pathway enrichment analysis of the genes (including *Gfi1*, **c**) with decreased mRNA levels in LSK from aged but not young Tet2 catalytic mutation (Mut) and Tet2 knockout (KO) mice in RNA-seq analysis (GSE227977). **d**. DNA methylation levels across *Gfi1* gene locus in WGBS analysis of HSPCs from young Tet2-mut, Tet2-KO and the wild-type control mice (GSE227977). Red box shows DMR. **e-g**. MeDIP-qPCR, RT-qPCR and western blot analysis of DNA methylation levels in *Gfi1*-specific DMR, mRNA and protein level of Gfi1 in Lin^-^ HSPCs from control and Tet2-KO young and aged mice. **h-j**. Colony formation analysis of the serial replating assay of the control and lentivirus-mediated Gfi1-overexpressed (OE) Lin^-^ HSPCs from aged Tet2-KO mouse. Data were normalized by input DNA (**e**) or β-actin (**f**), and compared with control groups. Data are the mean ± s.d. (**e, f, j**), two-tailed unpaired Student’s *t*-test. *, *P* < 0.05; **, *P* < 0.01; ***, *P* < 0.001

We further experimentally validated the increased DNA methylation level at *Gfi1* promoter in Tet2-KO Lin^-^ cells especially from aged mice (**Fig. 6e**), and decreased both mRNA and protein levels in *Tet2*-KO Lin^-^ cells from aged mice (**Fig. 6f, g**). We then observed the effect of rescuing Gfi1 expression in aged *Tet2*-KO HSCs, and found that overexpression of Gfi1 significantly reduced the replating potential of Tet2-KO Lin^-^ HSPCs from aged mice *in vitro* (**Fig. 6h-j**). These results further indicate that reduced expression of Gfi1 due to Tet2 loss-caused DNA hypermethylation contributes to MDS development at stem cell level.

## Discussion

Through establishing single-base resolution DNA methylomes of bone marrow primitive HSCs from healthy donors and MDS patients, we revealed the contribution of aberrant DNA methylation to HSC dysfunction in MDS at both single-gene locus and whole-genome levels. In the perspective of genome-wide DNA methylation variation in MDS, while hypermethylation was associated with regulatory regions of genes, hypomethylation was found in intergenic regions, including repeat elements. These intergenic hypomethylation may be inherited by offspring of MDS HSCs and the progenitors, which may increase genomic variation frequency to accelerate MDS development. It is worthwhile to note that tRNA genes are associated with hypermethylation in MDS HSCs, indicating that decreased synthesis of some proteins in HSCs may be a pathogenic factor for MDS. Moreover, since CD34^+^ bone marrow cells, which were used for DNA methylation analysis by most of the previous studies, cannot discriminate primitive HSCs from both myeloid progenitors and blast, our study provides valuable data for exploring how DNA methylation variations in primitive HSCs contribute to MDS development. Notably, DNA methylation variation in M3 case is different from those in M1 and M2 cases, indicating that the DNA methylation variations caused by heterogeneity among MDS patients in either the same subtype or among different MDS subtypes should be considered. Moreover, because of the small sample size of our study, future studies are needed to further observe the correlation of our identified DNA methylation variations, gene expression and clinical features of MDS cases like blast number and age.

To search for aberrant DNA methylation dysregulating genes that are potentially the pathogenic factors or therapeutic targets for MDS, we genome-widely identified DMR-associated genes between normal and MDS HSCs and found that cancer-related pathways were most enriched by hypermethylated DMR-associated genes. Further exploring DNA methylation dysregulating genes in our DMR data may lead to discover more cancer-related genes to provide potential biomarkers and targets for the treatment of cancers, especially for the leukemia transformed from MDS.

Some extrinsic pathways and intrinsic regulators regulating the self-renewal and differentiation of HSCs have been found to be associated with DMRs. For extrinsic pathways, besides these reported ones that are important for HSC maintenance, some cytokine-related pathways such as inflammatory and myeloid-specific ones were enriched by down-methylated DMR-associated genes. Dysregulation of these cytokines and their signal transducers may form an abnormal microenvironment to cause abnormality of bone marrow HSCs in MDS patients. For intrinsic regulators, epigenetic regulators and transcription factors with DNA methylation variations in at least two MDS samples are or may be involved in MDS development, e.g., somatic mutations and chromosomal rearrangements involving *RUNX1* are frequently observed in MDS^29^; Activation of HoxB4 permits long-term maintenance of functional hematopoietic stem and progenitor cells^30^; Mutation or suppression of the tumor suppressor mixed lineage leukemia 3 (MLL3) promotes self-renewal capacity of HSCs and impairs the differentiation of HSPCs^31,32^; HDAC5 inhibits HSC homing and engraftment^33^. Future studies are needed to investigate how aberrant DNA methylation dysregulates expression of these genes and their contribution to MDS development at HSC level.

Moreover, we focused on DNA methylation abnormalities in *GFI1* and *BMI1* gene loci, which were verified in other MDS samples. Zinc-finger repressor GFI1 was reported to restrict proliferation of HSCs and preserve HSC functional integrity. Loss of GFI1 leads to elevated proliferation rates, compromised repopulation function of HSC^34-36^. GFI1 is also import for differentiation and function of myeloid cell. Loss of GFI1 leads to severely neutropenic, accumulation of immature monocytic cells, and enhanced inflammatory immune response^37,38^. Reduced expression GFI1 leads to a fatal myeloproliferative disease (MPN) in *Gfi1*-knockdown mouse model^39^. GFI1-36N, which encodes a variant GFI1 protein with a decreased efficiency to act as a transcriptional repressor, was found to be clearly correlated with an increased risk of MDS patients to develop AML^40,41^. Expression of GFI1 was also reported to be repressed in MDS samples and low expression of GFI1 is associated with an inferior prognosis of AML patients^42^. For the epigenetic mechanism, GFI1 interacts with CoREST complex including histone deacetylase 1/2 (HDAC1/2) and lysine specific demethylase 1 (LSD1), which mediates transcriptional repression by GFI1^43^. Thus, our data provide an epigenetic mechanism for repressed expression of GFI1 during MDS development, implying that GFI1-dependent CoREST complex-mediated gene silencing is inhibited by DNA methylation in MDS HSCs. The target genes under the control this epigenetic regulation axis needs further investigation. As a component of Polycomb repressive complex 1 (PRC1) complex, BMI1 is essential for generating self-renewing adult HSCs and is required for efficient self-renewal of HSCs and leukemic stem cells^44,45^. Therefore, silencing GFI1 and overexpressing BMI1 by DNA methylation abnormalities in HSCs during MDS development may break the balance of quiescence and self-renewal of HSCs, contributing to transformation of hematopoietic progenitors.

TET2 is a well-established tumor suppressor for myeloid malignancies. The epigenetic mechanism underlying the role of TET2 in restricting transformation of HSCs is given serious attention. Using both a murine MDS and TET2-knockout model, we demonstrated that TET2 promoted the expression of GFI1 through promoter DNA demethylation in both MDS and aged HSCs. Repressed expression or impaired enzymatic activity of TET2, due to gene mutation or negative regulation mechanisms^46,47^, leads to promoter hypermethylation-mediated silencing of GFI1 expression, contributing to clonal hematopoiesis during MDS development and HSC aging. Dysregulation of Tet2-GFI1 epigenetic axis is a pathogenic mechanism for age-related hematopoietic diseases, including MDS, as well as myeloproliferative disorders, and myeloid leukemia^28^. Moreover, our study also reveals that loss of Tet2 in HSCs prevents the increase in expression of genes which are enriched in pathways of cancer, insulin resistance and neural signaling during aging. These genes may play repressive roles in the development of aging-related diseases, such as Gfi1 in repressing development of MDS and AML at stem cell level. Tet2 was recently reported to oxidize RNA m5C and antagonize MBD6-dependent H2AK119ub deubiquitination to repress aberrant expansion of myeloid leukemia cells^48^. Loss of Tet2 further accelerates MDS development through promoting the occurrence of secondary mutations in NHD13 mouse model^26^. Our study provides a DNA demethylation-dependent transcription regulation function of Tet2 in repressing myeloid cancers, especially during aging. However, more evidences from both *in vivo* experiments using Tet2-KO NHD13 mouse and clinical samples with Tet2 mutations are still needed for further validating our finding.

Notably, the region of TET2-mediated DNA demethylation in GFI1 gene locus was not the region significantly de novo methylated in human MDS HSCs compared with normal HSCs, implying that there is another mechanism in inhibiting DNA hypermethylation in HSCs beyond TET2 during MDS development. This mechanism is waiting to be revealed. For silencing GFI1 expression in MDS HSCs, TET2 loss acts as an accelerating role in further silencing GFI1. Thus, Tet2 deficiency can further expand hematopoietic stem and progenitor cell pool and accelerates leukemia transformation, further contributing to MDS development. Our study provides a mechanism in regulating DNA methylation for the pathogenic role of either loss-of-function mutations or expression and enzymatic activity inhibition of TET2 in MDS patients.

## Acknowledgment

We thank BGI for the assistance in bioinformatic analysis.

## Author contributions

X.C., Q.Z., Y.H. and L.H. conceived the study and designed the experiments; Y.H. and Y.G. sorted the HSCs from human BM samples; L.H., Y.G., Y.L., N.L. and B.Z. selected and characterized samples, provided disease-specific expertise in data analysis; J.L. provided the NHD13 transgenic mouse; L.H. and Q.S. performed experiments; Q.Z. conducted the bioinformatic analysis; X.C., Q.Z. and Y.H. analyzed the data and wrote the paper.

## Data availability

The Data for DNA methylomes of MDS and normal HSCs have been deposited into GEO. The accession numbers will be provided before publication. All other study data are included in the article and Supplemental information. All data are available in the main text or the supplementary materials.

## Materials and methods

### Human bone marrow collection and sample preparation

Bone marrow aspirates of 15 MDS patients with only supportive care and 10 healthy donors were collected between 2010 and 2011 and grouped according to the World Health Organization (WHO) classification system of MDS. The study was approved by the ethics committees of the Chinese People’s Liberation Army General Hospital and informed patient consent was obtained. All the bone marrow samples were harvested freshly and processed for flow sorting immediately. Mononuclear cells (MNCs) from the bone marrow samples were isolated by the Ficoll-Paque (GE Healthcare, Uppsala, Sweden) density gradient centrifugation. For purification of HSCs, MNCs were stained with anti-human lineage markers, CD34, CD133 (BioLegend, CA, USA), and a population of lin^-^CD34^+^CD133^+^ cells were sorted by MoFlo XDP cell sorter (Beckman-Coulter, CA, USA). The > 95% purity of the population was confirmed by performing FACS analysis of the sorted cells.

### Isolation of genomic DNA and bisulfite high-throughput sequencing

Total DNA was isolated using a QIAamp DNA Mini Kit (QIAGEN, Hilden, Germany) following the manufacturer’s instructions. Bisulfite DNA was prepared and high-throughput sequencing was performed as previously described^20^.

### Reverse Transcription qPCR (RT-qPCR)

Reverse transcription products using a reverse transcription system (Toyobo) was used for quantitative PCR (qPCR) analysis by LightCycler (Roche) and SYBR RT-PCR kit (Takara) as described previously^11^. Data were normalized by the level of β-actin.

### Bisulfite sequencing alignment and identification of methyl-cytosines

The bisulfite treated reads generated by Illumina sequencing were aligned to Hg18, and methyl-cytosine identification was performed as previously reported^20^.

### MeDIP assay

Genomic DNA was extracted using QIAamp DNA mini kit (QIAGEN) and fragmented into a main band at between 200bp and 400bp using the Bioruptor from Diagenode. The purified genomic DNA was denatured at 95°C for 10 min and incubated with 5-mC antibody (Zymo Research) at 4°C overnight. Protein A/G magnetic beads were added to pull-down antibody-DNA complexes for 2 hr at 4°C. Beads were washed and digested by protease K. Eluted DNA was extracted with phenol-chloroform, ethanol precipitated. Obtained DNA from MeDIP assay was subjected to qPCR analysis.

### Identification of differentially methylated regions (DMRs)

All the shared cytosines (Cs) in CG context which had coverage over 1× in each of the six samples were selected for DMR analysis. DMRs were identified by sliding windows each of which contains 5 CGs with the following steps: Step 1, windows which have a mean depth over 10 and 80% CGs coverage were used; Step 2, the window with the DNA methylation rate being at least double difference (Fisher test, p=<0.01) and either of the DNA methylation level being higher than 50% were identified as differentially methylated window; Step 3, each identified differentially methylated window was extended unless two CGs were at a distance of over 200bp or it no longer meet above standards for differentially methylated window. The extended window was the DMR.

DMRs between each normal and disease sample were firstly searched. But if DNA methylation variation of the DMR was also significant among three normal samples for 3×2 fisher test, the DMR was discarded. If the DMR appeared spontaneously in two or three MDS samples, it was used for the analysis. The gene (from upstream 10Kb to transcription end site (TES)) with at least one DMR was defined as DMR-associated gene.

### DNA methylation pattern discrepancy between MDS and normal samples

To investigate DNA methylation pattern discrepancy between MDS and normal samples, DNA methylation level was computed as the average of it for every Cs in CG or non-CG context in genomic element. Alternatively, bigwig files of DNA methylation levels in CG context were used for PCA and DMR-based heatmap analysis using deepTools.

### Pathway enrichment analysis

DMRs-associated genes were subjected to pathway enrichment analysis based on DAVID database.

### Mice

*Tet2*-knockout mice with a C57BL/6×129/SvEv background were provided by Dr. R. L. Levine. NUP98-HOXD13 (NHD13) transgenic mice were obtained from the Jackson Laboratory^49^. All mice were maintained under specific pathogen-free conditions. All animal experiments were performed according to National Institutes of Health Guide for the Care and Use of Laboratory Animals, with the approval of the Scientific Investigation Board of Naval Medical University, Shanghai, China.

### Lentivirus-mediated gene knockdown or overexpression

Short hairpin RNAs (shRNAs) specifically targeting to Tet2 were generated by GenePharma (Shanghai, China). Lin^-^ HSPC were isolated from bone marrow of Tet2-KO or NHD13 mice using EasySep™ negative selection kit (STEMCELL Technologies) and infected with lentivirus expression Tet2-specific shRNA, Gfi1, or controls. Cells were infected with a multiplicity of infection (MOI) of 50 for 4 days in SFEM medium (#09650, Stem Cell Technologies), or Methyl-cellulose Media (M3434; STEMCELL Technologies) with myeloid differentiation cytokines.

### Colony-forming assays

Lin^-^ HSPC were isolated from bone marrow of Tet2-KO or NHD13 mice, infected with lentivirus for Tet2 silencing or Gfi1 overexpression were seeded into Methyl-cellulose Media (M3434; STEMCELL Technologies) with myeloid differentiation cytokines. After incubation for 7 days, colonies were counted, or were then sequentially replated another 7 days for replating assay.

### Statistical analysis

Error bars displayed throughout the paper represent s.d. and were calculated from triplicate technical. Sample sizes were chosen by standard methods to ensure adequate power, and no randomization of weight and sex or blinding was used for animal studies. Data shown are representative of three independent experiments. No statistical method was used to predetermine sample size. Statistical significance was determined using unpaired two-sided Student’s *t*-test, with *P* < 0.05 considered statistically significant.

## References

1 Raskovalova, T., Jacob, M. C. & Park, S. Myelodysplastic Syndromes. N Engl J Med 383, 2590 (2020).

2 Will, B. et al. Stem and progenitor cells in myelodysplastic syndromes show aberrant stage-specific expansion and harbor genetic and epigenetic alterations. Blood 120, 2076–2086 (2012).

3 Meng, Y. & Nerlov, C. Epigenetic regulation of hematopoietic stem cell fate. Trends Cell Biol 35, 217–229 (2025).

4 Cheng, H., Zheng, Z. & Cheng, T. New paradigms on hematopoietic stem cell differentiation. Protein Cell 11, 34–44 (2020).

5 Figueroa, M. E. et al. MDS and secondary AML display unique patterns and abundance of aberrant DNA methylation. Blood 114, 3448–3458 (2009).

6 Thoms, J. A. I. et al. Clinical response to azacitidine in MDS is associated with distinct DNA methylation changes in HSPCs. Nat Commun 16, 4451 (2025).

7 del Rey, M. et al. Genome-wide profiling of methylation identifies novel targets with aberrant hypermethylation and reduced expression in low-risk myelodysplastic syndromes. Leukemia 27, 610–618 (2013).

8 Reilly, B. et al. DNA methylation identifies genetically and prognostically distinct subtypes of myelodysplastic syndromes. Blood Adv 3, 2845–2858 (2019).

9 Lopez-Moyado, I. F., Ko, M., Hogan, P. G. & Rao, A. TET Enzymes in the Immune System: From DNA Demethylation to Immunotherapy, Inflammation, and Cancer. Annu Rev Immunol 42, 455–488 (2024).

10 Cong, B., Zhang, Q. & Cao, X. The function and regulation of TET2 in innate immunity and inflammation. Protein Cell 12, 165–173 (2021).

11 Shen, Q. et al. Tet2 promotes pathogen infection-induced myelopoiesis through mRNA oxidation. Nature 554, 123–127 (2018).

12 Zhang, Q. et al. Tet2 is required to resolve inflammation by recruiting Hdac2 to specifically repress IL-6. Nature 525, 389–393 (2015).

13 Kohli, R. M. & Zhang, Y. TET enzymes, TDG and the dynamics of DNA demethylation. Nature 502, 472–479 (2013).

14 Moran-Crusio, K. et al. Tet2 loss leads to increased hematopoietic stem cell self-renewal and myeloid transformation. Cancer Cell 20, 11–24 (2011).

15 Cimmino, L. et al. Restoration of TET2 Function Blocks Aberrant Self-Renewal and Leukemia Progression. Cell 170, 1079–1095 e1020 (2017).

16 Ko, M. et al. Impaired hydroxylation of 5-methylcytosine in myeloid cancers with mutant TET2. Nature 468, 839–843 (2010).

17 Jiang, Y. et al. Aberrant DNA methylation is a dominant mechanism in MDS progression to AML. Blood 113, 1315–1325 (2009).

18 Sekeres, M. A. & Taylor, J. Diagnosis and Treatment of Myelodysplastic Syndromes: A Review. JAMA 328, 872–880 (2022).

19 Stomper, J., Rotondo, J. C., Greve, G. & Lubbert, M. Hypomethylating agents (HMA) for the treatment of acute myeloid leukemia and myelodysplastic syndromes: mechanisms of resistance and novel HMA-based therapies. Leukemia 35, 1873–1889 (2021).

20 Li, Y. et al. The DNA methylome of human peripheral blood mononuclear cells. PLoS Biol 8, e1000533 (2010).

21 Takahashi, M. et al. CD133 is a positive marker for a distinct class of primitive human cord blood-derived CD34-negative hematopoietic stem cells. Leukemia 28, 1308–1315 (2014).

22 Shastri, A., Will, B., Steidl, U. & Verma, A. Stem and progenitor cell alterations in myelodysplastic syndromes. Blood 129, 1586–1594 (2017).

23 Smith, Z. D., Hetzel, S. & Meissner, A. DNA methylation in mammalian development and disease. Nat Rev Genet 26, 7–30 (2025).

24 Papanicolau-Sengos, A. & Aldape, K. DNA Methylation Profiling: An Emerging Paradigm for Cancer Diagnosis. Annu Rev Pathol 17, 295–321 (2022).

25 de Jong, M. M. E., Chen, L., Raaijmakers, M. & Cupedo, T. Bone marrow inflammation in haematological malignancies. Nat Rev Immunol 24, 543–558 (2024).

26 Huang, F. et al. TET2 deficiency promotes MDS-associated leukemogenesis. Blood Cancer J 12, 141 (2022).

27 Chen, B. Y. et al. SETD2 deficiency accelerates MDS-associated leukemogenesis via S100a9 in NHD13 mice and predicts poor prognosis in MDS. Blood 135, 2271–2285 (2020).

28 Flores, J. C. et al. Comparative analysis of Tet2 catalytic-deficient and knockout bone marrow over time. Exp Hematol 124, 45–55 e42 (2023).

29 Sood, R., Kamikubo, Y. & Liu, P. Role of RUNX1 in hematological malignancies. Blood 129, 2070–2082 (2017).

30 Izawa, K. et al. Activated HoxB4-induced hematopoietic stem cells from murine pluripotent stem cells via long-term programming. Exp Hematol 89, 68–79 e67 (2020).

31 Chen, C. et al. MLL3 is a haploinsufficient 7q tumor suppressor in acute myeloid leukemia. Cancer Cell 25, 652–665 (2014).

32 Chen, R. et al. Kmt2c mutations enhance HSC self-renewal capacity and convey a selective advantage after chemotherapy. Cell Rep 34, 108751 (2021).

33 Huang, X., Guo, B., Liu, S., Wan, J. & Broxmeyer, H. E. Neutralizing negative epigenetic regulation by HDAC5 enhances human haematopoietic stem cell homing and engraftment. Nat Commun 9, 2741 (2018).

34 Gilks, C. B., Bear, S. E., Grimes, H. L. & Tsichlis, P. N. Progression of interleukin-2 (IL-2)-dependent rat T cell lymphoma lines to IL-2-independent growth following activation of a gene (Gfi-1) encoding a novel zinc finger protein. Mol Cell Biol 13, 1759–1768 (1993).

35 Zeng, H., Yucel, R., Kosan, C., Klein-Hitpass, L. & Moroy, T. Transcription factor Gfi1 regulates self-renewal and engraftment of hematopoietic stem cells. EMBO J 23, 4116–4125 (2004).

36 Hock, H. et al. Gfi-1 restricts proliferation and preserves functional integrity of haematopoietic stem cells. Nature 431, 1002–1007 (2004).

37 Karsunky, H. et al. Inflammatory reactions and severe neutropenia in mice lacking the transcriptional repressor Gfi1. Nat Genet 30, 295–300 (2002).

38 Hock, H. et al. Intrinsic requirement for zinc finger transcription factor Gfi-1 in neutrophil differentiation. Immunity 18, 109–120 (2003).

39 Fraszczak, J. et al. Reduced expression but not deficiency of GFI1 causes a fatal myeloproliferative disease in mice. Leukemia 33, 110–121 (2019).

40 Botezatu, L. et al. GFI1(36N) as a therapeutic and prognostic marker for myelodysplastic syndrome. Exp Hematol 44, 590–595 e591 (2016).

41 Khandanpour, C. et al. A variant allele of Growth Factor Independence 1 (GFI1) is associated with acute myeloid leukemia. Blood 115, 2462–2472 (2010).

42 Moroy, T. & Khandanpour, C. Role of GFI1 in Epigenetic Regulation of MDS and AML Pathogenesis: Mechanisms and Therapeutic Implications. Front Oncol 9, 824 (2019).

43 Saleque, S., Kim, J., Rooke, H. M. & Orkin, S. H. Epigenetic regulation of hematopoietic differentiation by Gfi-1 and Gfi-1b is mediated by the cofactors CoREST and LSD1. Mol Cell 27, 562–572 (2007).

44 Smith, L. L. et al. Functional crosstalk between Bmi1 and MLL/Hoxa9 axis in establishment of normal hematopoietic and leukemic stem cells. Cell Stem Cell 8, 649–662 (2011).

45 Yamagishi, M., Iwama, A. & Kitabayashi, I. Polycomb Repressive Complexes as Therapeutic Targets in Hematologic Malignancies. Exp Hematol, 105339 (2025).

46 Ogawa, S. Genetics of MDS. Blood 133, 1049–1059 (2019).

47 Sun, J. et al. SIRT1 Activation Disrupts Maintenance of Myelodysplastic Syndrome Stem and Progenitor Cells by Restoring TET2 Function. Cell Stem Cell 23, 355–369 e359 (2018).

48 Zou, Z. et al. RNA m(5)C oxidation by TET2 regulates chromatin state and leukaemogenesis. Nature 634, 986–994 (2024).

49 Xu, H. et al. Single-cell transcriptome sequencing reveals the mechanism of Realgar improvement on erythropoiesis in mice with myelodysplastic syndrome. Cancer Cell Int 25, 135 (2025).

