## Supplementary materials for "Base-resolution DNA methylome of human MDS hematopoietic stem cell reveals TET2-GFI1 epigenetic axis repressing MDS"

**Supplementary Table 1, 3, 4 and Supplementary Figure 1-5**

**Table S1 General characteristics of MDS patients** and healthy donors

| **characteristics**  **N** | **sex,M/F** | **Age, average**  **(y±SD)** | **Blast,%**  **（y±SD）** | **Cytogenetics, normal/abnormal** |
| --- | --- | --- | --- | --- |

| RAEB1/2  for sequencing | 3 | 2/1 | 24±11 | 9±2 | 0/3 |
| --- | --- | --- | --- | --- | --- |
| RAEB1/2  for verification | 9 | 9/0 | 39±13 | 10±8 | 3/6 |
| tAML  for verification | 2 | 2/0 | 21±3 | 28±4 | 1/1 |
| RA  for verification | 1 | 1/0 | 35 | 1 | 1/0 |
| normal  for sequencing | 3 | 2/1 | 29±5 | - | - |
| normal  for verification | 7 | 4/3 | 33±11 | - | - |

**Table S3 Pathway enrichment analysis of DMR-**associated genes

| **Category** | **Term** | **P Value** |
| --- | --- | --- |
| **Up-methylated DMR**  KEGG_PATHWAY | Pathways in cancer | 3.84E-6 |
| KEGG_PATHWAY | Basal cell carcinoma | 7.42E-4 |
| BIOCARTA | Arf Inhibits Ribosomal Biogenesis | 0.00106 |
| KEGG_PATHWAY | Melanogenesis | 0.00166 |
| KEGG_PATHWAY | Melanoma | 0.00461 |
| KEGG_PATHWAY | Tight junction | 0.00770 |
| KEGG_PATHWAY | MAPK signaling pathway | 0.00961 |
| KEGG_PATHWAY | Wnt signaling pathway | 0.01968 |
| KEGG_PATHWAY | Maturity onset diabetes of the young | 0.02510 |
| BIOCARTA | Transcription Regulation by CARM1 | 0.03474 |
| KEGG_PATHWAY | Hedgehog signaling pathway | 0.04088 |
| KEGG_PATHWAY | Neuroactive ligand-receptor interaction | 0.04320 |
| BIOCARTA | NFAT and Hypertrophy of the heart | 0.04966 |
| **Down-methylated DMR** |  |  |
| KEGG_PATHWAY | Jak-STAT signaling pathway | 0.00474 |
| KEGG_PATHWAY | Natural killer cell mediated cytotoxicity | 0.00622 |
| BIOCARTA | Erythrocyte Differentiation Pathway | 0.03216 |
| KEGG_PATHWAY | Regulation of actin cytoskeleton | 0.03439 |
| BIOCARTA | Regulation of hematopoiesis by cytokines | 0.04212 |
| KEGG_PATHWAY | Cytokine-cytokine receptor interaction | 0.04532 |
| KEGG_PATHWAY | Aldosterone-regulated sodium reabsorption | 0.05300 |
| KEGG_PATHWAY | Insulin signaling pathway | 0.06174 |
| KEGG_PATHWAY | Pathways in cancer | 0.07826 |
| BIOCARTA | CXCR4 Signaling Pathway | 0.09093 |

**Table S4 Functional classification of DMR-associated genes based on** KEGG database

| Functional class | Gene number |
| --- | --- |
| Enzymes | 522 |
| Transcription factors | 207 |
| Cell adhesion molecules and their ligands | 149 |
| Protein kinases | 106 |
| Chromosome | 103 |
| Cellular antigens | 87 |
| Transporters | 86 |
| Ubiquitin system | 79 |
| Ion channels | 75 |
| Peptidases | 69 |
| Solute carrier family | 65 |
| G protein-coupled receptors | 64 |
| Cytoskeleton proteins | 62 |
| Cytokine receptors | 49 |
| Glycosyltransferases | 43 |
| Cytokines | 43 |
| Heparan sulfate/heparin binding proteins | 42 |
| Enzyme-linked receptors | 32 |
| Spliceosome | 31 |
| GTP-binding proteins | 28 |
| Transcription machinery | 22 |
| DNA repair and recombination proteins | 22 |
| Glycan binding proteins | 17 |
| Chaperones and folding catalysts | 14 |
| DNA replication proteins | 14 |
| Proteoglycans | 12 |
| Lipid biosynthesis proteins | 12 |

**Supplementary Figures:**

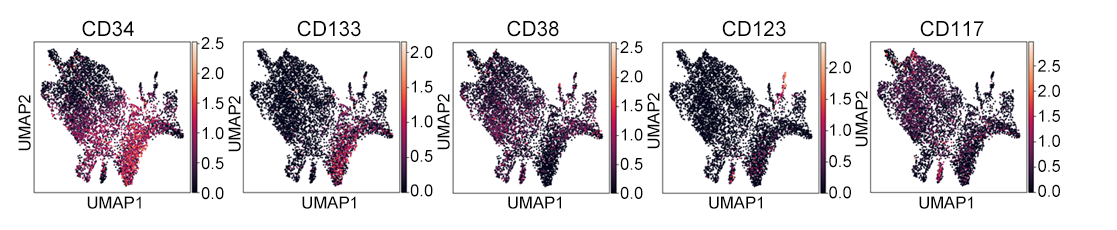

**Fig. S1 Human Lin^-^CD34^+^CD133^+^ cells are primitive HSCs**

Dot plot analysis of expression of indicated genes in scRNA-seq data of bone marrow CD34^+^ cells from MDS patients (GSM7842961).

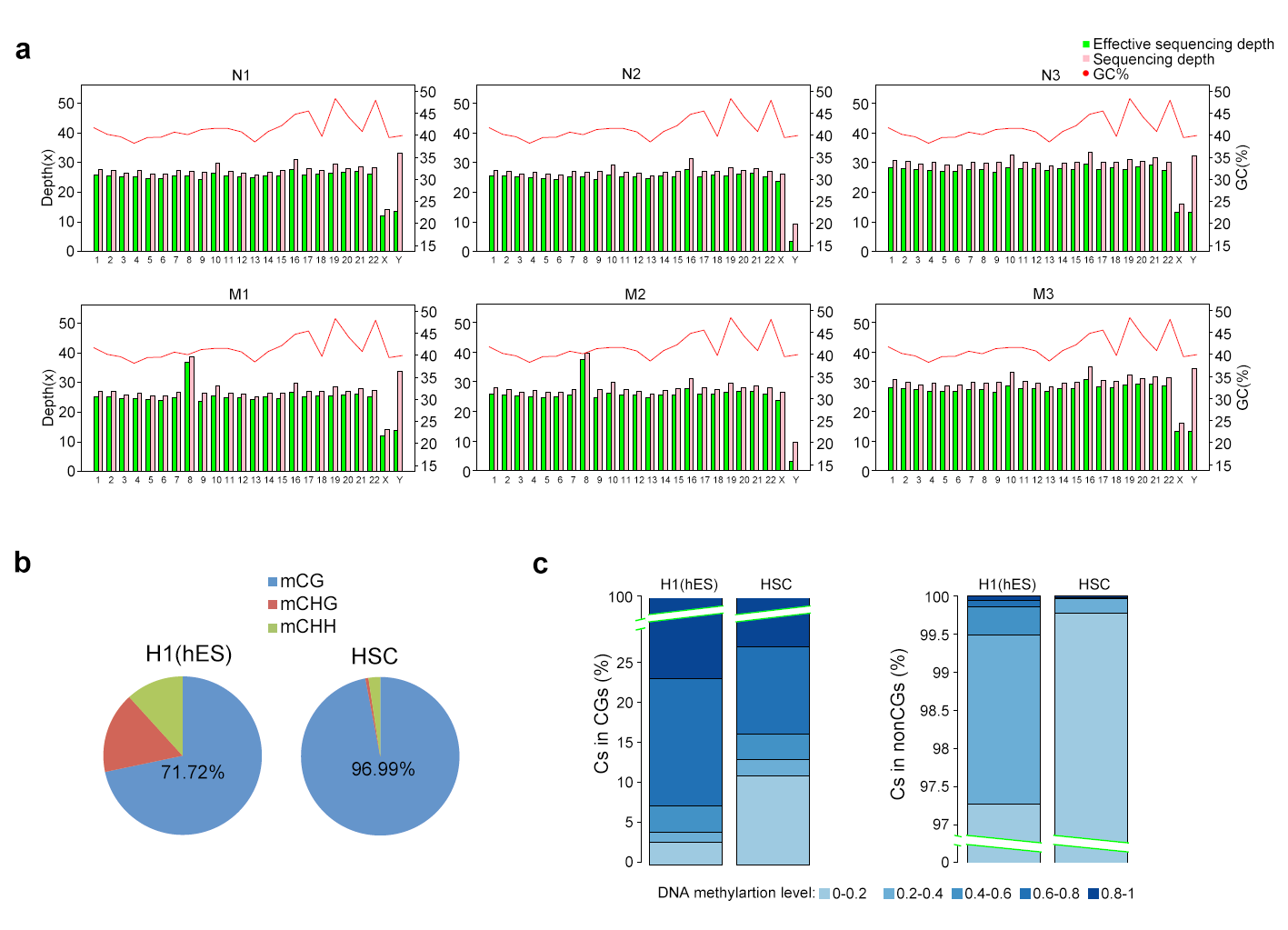
**Fig. S2 DNA methylome data and patterns in normal and MDS HSCs**

**a.** Effective sequencing depth referred to unique mapped reads.

**b.** The proportion of methylcytosines (mCs) in CG, CHG, and CHH sequence context in the hES and normal HSCs (combined data of normal controls). H represents any nucleotide except G.

**c.** Distribution of DNA methylation levels of cytosines (Cs) of the corresponding sequence context in the indicated stem cells. The differently colored numbers represent the intervals of DNA methylation levels.

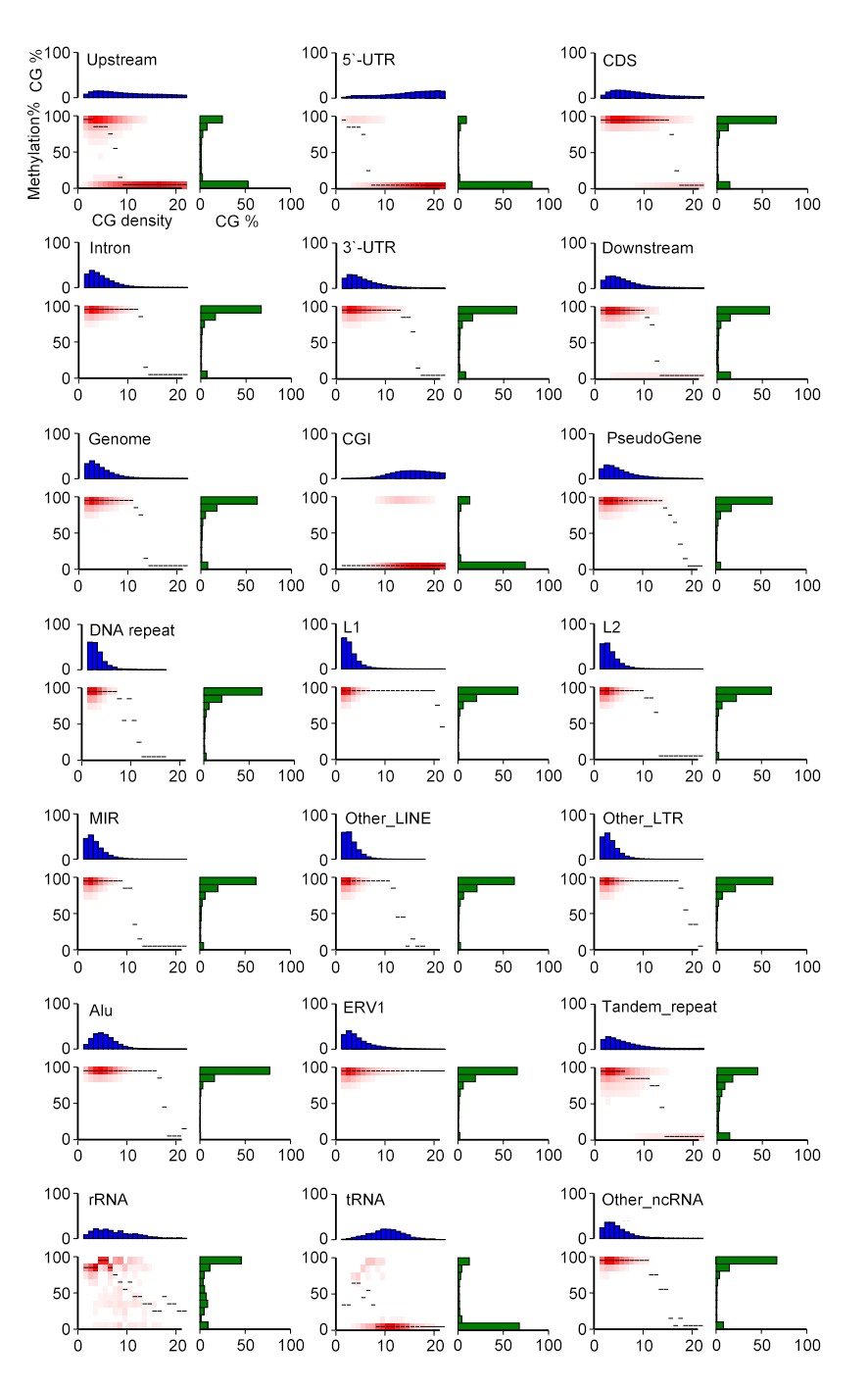

**Fig. S3 Heatmaps of distinct DNA methylation levels in CG context and CG densities in different genomic elements**

Shared Cs in CG context in three normal HSCs genomes (depth>=5) were analyzed. CG density (x-axis) was defined as number of CG dinucleotides in 200bp windows. DNA methylation level (y-axis) was defined as mean methylation level of Cs in CG context of the three samples. The thin black lines within each panel denoted the median methylation level of Cs at the given local density. The red color gradients indicated abundances of Cs that fell into bins of given DNA methylation levels and CG densities. The blue bar charts above heatmaps showed the distribution of CG densities, projected onto the x-axis of the heatmaps. The green bar charts to the right of the heatmaps showed the distribution of methylation levels, projected onto the y-axis of the heat maps.

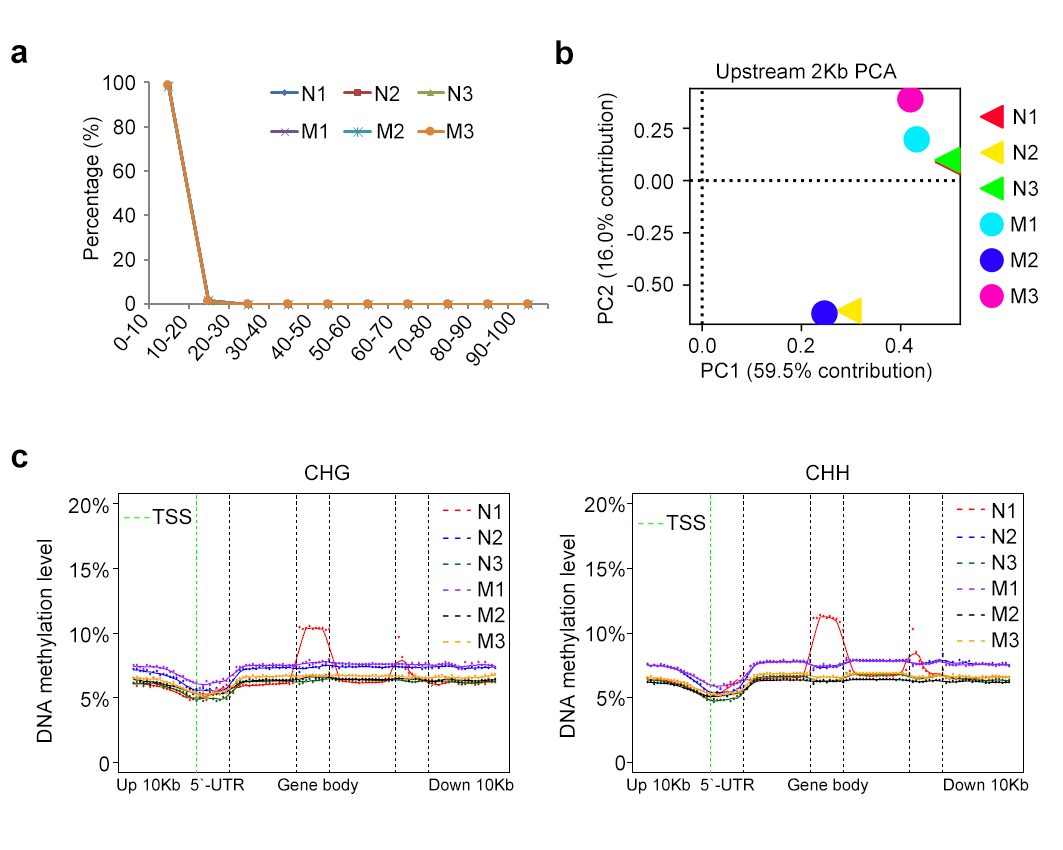

**Fig. S4 DNA methylation levels of MDS and normal HSCs**

**a.** Distribution of DNA methylation levels of cytosines (Cs) in non-CG context (CHG and CHH).

**b.** PCA analysis of DNA methylation levels of proximal promoters (upstream 2Kb) of normal and MDS HSCs.

**c.** Mean DNA methylation levels for cytosines (Cs) in non-CG context (CHG and CHH) with different features across all gene loci. Mean DNA methylation level of each equal-sized bin was calculated.

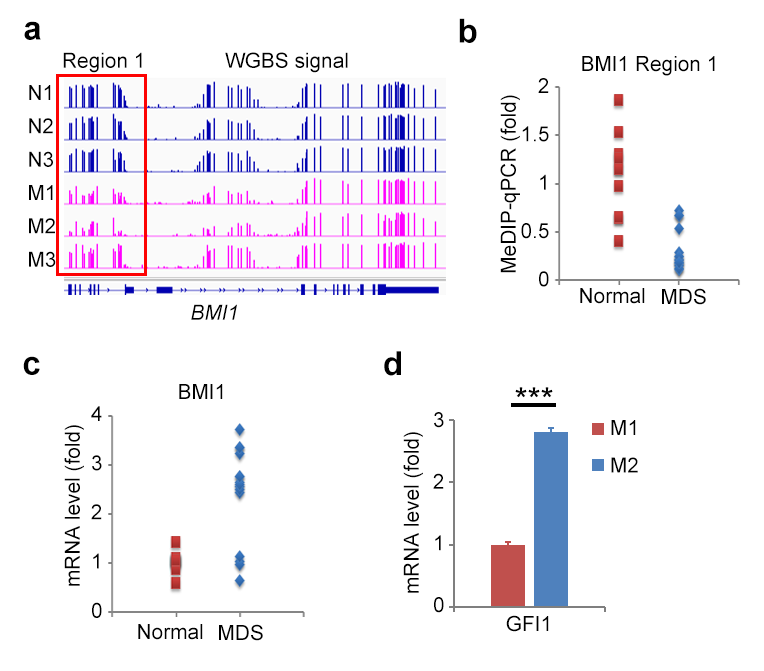

**Fig. S5 DNA demethylation increases BMI1 expression in MDS HSCs**

**a.** DNA methylation levels in CG context across *BMI1* genes in genome browser view. Red boxes show regions with DNA methylation variations in normal (N) and MDS (M) HSCs.

**b, c.** MeDIP-qPCR analysis of DNA methylation levels of region 1 in *BMI1* locus and RT-qPCR analysis of mRNA levels of BMI1 in normal (n = 10) and MDS (n = 15) HSCs.

**d.** RT-qPCR analysis of mRNA levels of GFI1 in HSCs of M1 and M2 MDS samples. Signals were normalized by β-actin.

Data were normalized by input DNA (**b**) or β-actin (**c**, **d**), and compared with control groups. Data are the mean ± s.d. (**d**), two-tailed unpaired Student’s *t*-test. *, *P* < 0.05; **, *P* < 0.01; ***, *P* < 0.001

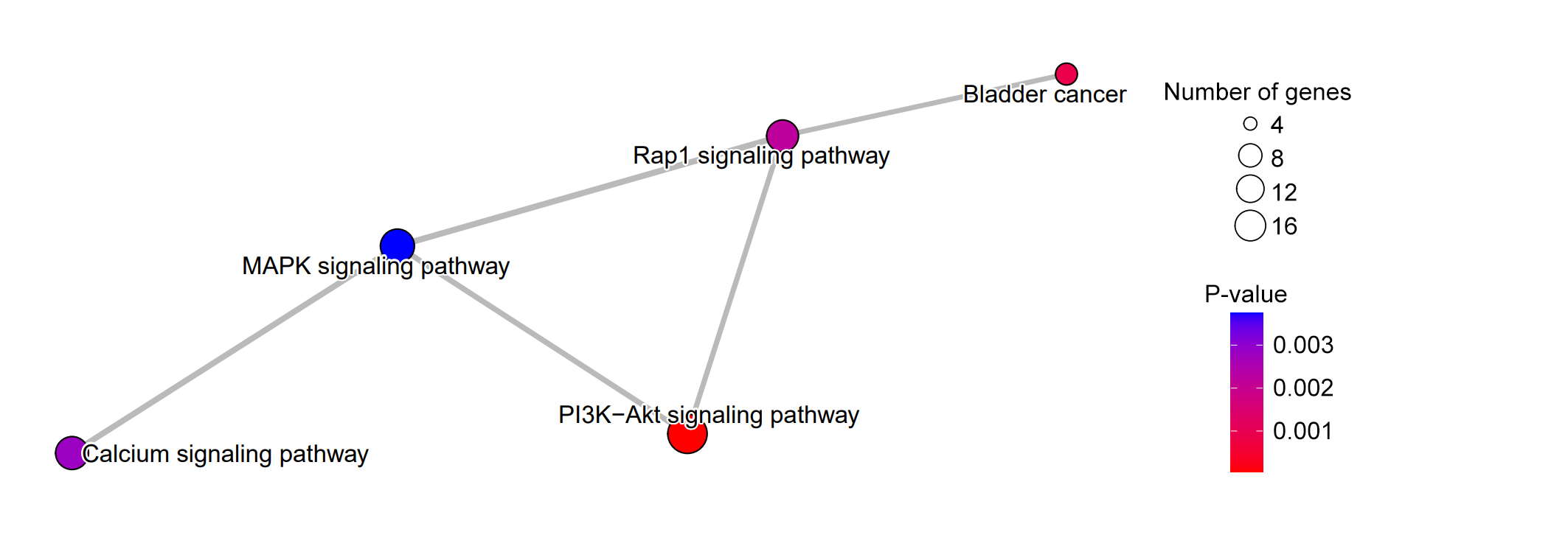

**Fig. S6 Interaction map of KEGG pathways**

The sizes and colors of the circles respectively indicate the number and enrichment significance of genes in the indicated pathways.
